# Death in the taste bud: engulfment of dying taste receptor cells by glial-like Type I cells

**DOI:** 10.1101/2024.09.06.611711

**Authors:** Courtney E. Wilson, Robert S. Lasher, Ernesto Salcedo, Ruibiao Yang, Yannick Dzowo, John C. Kinnamon, Thomas E. Finger

## Abstract

Taste buds comprise 50-100 epithelial derived cells, including glial-like cells (Type I) and two types of receptor cells (Types II and III). All of these taste cells are renewed throughout the life of an organism from a pool of uncommitted basal cells. Immature cells enter the bud at its base, maturing into one of the three mature cell types. How taste cells die and/or exit the bud, as well as the role of the glial-like cells in this process, remains unclear. Here we present morphological data obtained through Serial Blockface Scanning Electron Microscopy of murine circumvallate taste buds, revealing taste cells at the end of their life. Cells we identify as dying share morpholowgical features typical of apoptosis: swollen endoplasmic reticulum, large lysosomes, degrading organelles, distended outer nuclear membranes, heterochromatin reorganization, cell shrinkage, and cell and/or nuclear fragmentation. Most early stage dying cells have Type II cell morphologies, while a few display Type III cell features. Many dying cells maintain contacts with nerve fibers, but those postsynaptic fibers often appear to be detached from the main trunk of an afferent nerve. Dying cells, like mature Type II and III taste cells, are surrounded by glial-like Type I cells. In many instances Type I cells appear to be engulfing their dying neighbors, suggesting for them a novel, phagocytic role. Surprisingly, virtually no Type I cells display features of apoptosis, although they reportedly have the shortest residence time in taste buds. The ultimate fate of Type I cells therefore remains unknown.

**Table of Contents Entry:** *Main Points:*

1. Dying taste cells display morphologies consistent with apoptosis.
2. Glial-like Type I cells engulf dying neighbors, perhaps acting as “undertakers.”
3. We find no dying Type I cells—all early dying cells are taste transducing Type II or III cells.

## Introduction

The peripheral sensory organs for hearing, balance, and taste all contain glial-like cell populations as well as secondary sensory cells involved directly in transduction. These glial-like cells share many features with astrocytes in the CNS including: uptake and inactivation of neurotransmitters (e.g. expression of GLAST, ectoNTPDase) (Furness and Lehre 1997; Glowatzki et al., 2006; Vlajkovic et al., 2002), regulation of ionic environment (K uptake and extrusion) (Li et al., 2012), and metabolic and structural support. Many of these molecular characteristics are also shared by the Muller glial cells of the retina (Bringmann et al., 2009; Wang et al., 2017). In taste buds, the glial-like cells are identified as Type I cells in contradistinction to the Type II and Type III cells involved in transduction of potential taste substances. Type I cells express GLAST (Lawton et al., 2000) and ectoATPase (Bartel et al., 2006), and are implicated in the regulation of potassium ion flux and extracellular neurotransmitter levels (Dvoryanchikov et al., 2009; Rodriguez et al., 2021). Unlike the glial-like cells of other sensory organs, Type I taste cells are thought to have a short life span, being repeatedly replaced over the span of several weeks like the other cells in the bud (Farbman, 1980; Perea-Martinez et al., 2013).

To maintain a stable population, taste buds must lose and gain cells at roughly the same rate. Organisms rid themselves of aging or unhealthy cells by three general mechanisms: apoptosis, autophagy, and necrosis (D’Arcy, 2019). Apoptosis is a form of programmed cell death resulting in characteristic ultrastructural changes: chromatin condensation, nuclear fragmentation, cell shrinkage, and ultimately the formation of apoptotic bodies that fragment off the cell. In contrast, in autophagy, which can either protect or kill a cell, macroproteins and organelles are sequestered into autophagosomes and are degraded. The process of necrosis may result from acute injury to the cell; necrotic cells swell and rupture, spilling their contents and triggering an inflammatory response (D’Arcy, 2019; Chen et al., 2020). Only a few studies of cell death in the taste bud exist, and all report instances of taste cell apoptosis (Suzuki et al., 1996A; Takeda et al., 1996; Zeng and Oakley, 1999; Zeng et al., 2000; Huang and Lu, 2001; Ueda et al., 2008).

We examined serial electron micrographs through taste buds obtained by serial blockface scanning electron microscopy (sbfSEM) to identify ultrastructural signs of cell death, expecting to find dying glial-like Type I cells as well as taste-transducing Type II and III cells in proportion to their respective populations and rates of replacement within taste buds. The three mature taste cell types are each unique in morphology, lifespan, and function. Glial-like Type I cells constitute ∼50% of the mature cells in the bud (Murray et al., 1969; Pumplin et al., 1997; Yang et al., 2020) and have an estimated lifespan of ∼7 and ∼16 days in rat and mouse, respectively (Farbman, 1980; Perea-Martinez et al., 2013). Type II cells, about one quarter of the cells in a bud, are spindle-shaped receptor cells and live for ∼14-30 days (Perea-Martinez et al., 2013; Gross et al., 2017; Yang et al., 2020). Each Type II cell expresses receptors for only one of the classical taste qualities: bitter, sweet, or umami, and perhaps amiloride-dependent sodium taste (Ohmoto et al., 2020). Type II cells communicate to nerve fibers via channel synapses, involving large-pore, voltage-gated channels associated with “atypical” mitochondria (Royer and Kinnamon, 1988; Chaudhari and Roper, 2010; Taruno et al., 2013; Romanov et al., 2018). The other receptor cell type of taste buds, Type III cells, are spindle-shaped cells, which constitute about 15-17% of the mature cells in a bud and are the most long-lived taste cells, with a half-life of at least 22 days (Perea-Martinez et al., 2013). Type III cells transduce ionic taste qualities, i.e. sour and highly salty stimuli, and communicate to nerves via conventional vesicular synapses (Kinnamon et al., 1985; Yee et al., 2001; Huang et al., 2008; Yang et al., 2020). We hypothesize that Type I cells, being the most abundant and shortest-lived cell type in taste buds, would exhibit the clearest signs of apoptosis. Surprisingly, we did not find any glial-like Type I cells with morphological signs of cell death in our datasets. Rather, dying Type II and III taste cells are all at least partially surrounded by Type I taste cells, some quite substantially so. This arrangement and the presence of abundant, large lysosomes in Type I cells adjacent to dying cells suggest that Type I cells, like astrocytes in the CNS (Morizawa et al., 2017; Kono et al., 2021; Zhou et al., 2022), can engulf and degrade dying cells. That we find no evidence of dying Type I cells leaves open the question as to when and how these cells exit the taste bud to be replaced during the lifespan.

## Methods

### Serial Block Face Scanning Electron Microscopy

As reported previously, Serial Blockface Scanning Electron Microscopy (sbfSEM) [see (Yang et al., 2020)] was used to generate two datasets from adult (>45 days) mouse circumvallate taste buds: DS2, composed of 563 sections, and TF21, composed of 633 sections. The data used in the present study were obtained primarily from two complete and one nearly complete taste buds from TF21 (Fig.1). A small number of cells from DS2 were also identified as early stage dying cells and more from this dataset were used for analysis of lysosomes. The original image data are freely available at the Electron Microscopy Public Image Archive, part of the European Bioinformatics Institute (EMBL-EBI), in dataset EMPIAR-10331. Every taste cell in each section was assigned a unique identifier and was analyzed using Reconstruct software (Synapse Web Reconstruct, RRID:SCR_002716) (Fiala, 2005) to determine cell type and generate 3D reconstructions.

**Figure 1.**
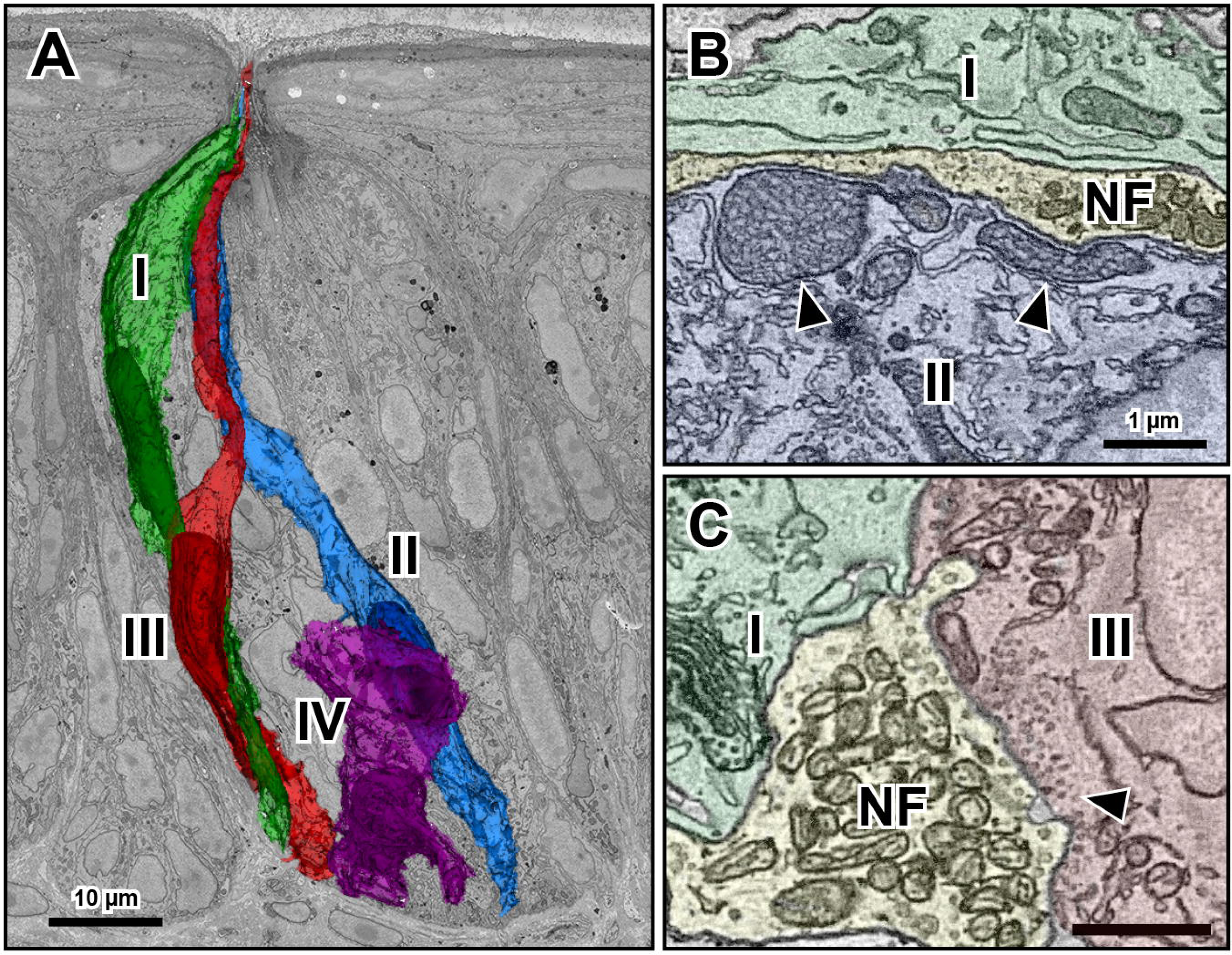
Taste cell types and synapses in murine circumvallate taste buds. **A.**Electron micrograph of a circumvallate mouse taste bud with overlaid individual reconstructions of a Type I cell (green), Type II cell (blue), Type III cell (red), and Type IV immature cell (purple). Scale bar is 10 µm. **B.** Micrograph of the channel synapse between a Type II cell (blue) and adjacent nerve fiber (yellow) bordered by a Type I cell (green). Arrows indicate the large, atypical mitochondria characteristic of Type II cell synapses. Scale bar is 1 µm for both **B** and **C**. **C.** Micrograph of the synapse between a Type III cell (red) and adjacent nerve fiber (yellow) bordered by a Type I cell (green). Arrow indicates the cluster of vesicles at the pre-synaptic membrane.

### Identifying taste cells

Mammalian taste buds contain three basic types of mature taste cells: Type I, II, and III cells. We identified cells as being part of one of these categories based on several morphological features described in previous studies (Yang et al., 2020) (**Figure 1**). Type I cells tend to have elongate, invaginated nuclei, as well as thin, lamellar processes that extend around and between neighboring taste cells. While they often wrap around and border innervating nerve fibers, they do not display synaptic morphology (either atypical mitochondria or synaptic vesicles) at these sites of contact. The apical structure of Type I cells falls into two main categories: bushy, with many short microvilli, or arboriform, with one main microvillus with smaller microvillar “branches” (Yang et al., 2020). Type II cells are elongate, have a somewhat flattened fusiform shape, and relatively smooth, ovoid nuclei. The majority of Type II cells contain so called “atypical” mitochondria. These large mitochondria are characterized by tubular rather than stacked cristae and exist at points of synaptic contact from Type II cells onto afferent nerve fibers (Royer and Kinnamon 1988; Romanov et al., 2018) (**Figure 1B**). At their apical regions, Type II cells (e.g., **Figure 2A**) generally feature a single, large microvillus that extends far into the taste pore. Type III cells are elongate and spindle-shaped, with a single, large microvillus that extends far into the taste pore and nuclei that are elongate and slightly invaginated (Yee et al., 2001; Yang et al., 2020). They are the only cells in the taste bud that form conventional, vesicular synapses onto afferent nerve fibers. These points of synaptic contact feature clear, 40-60nm diameter vesicles clustered at the plasma membrane region bordering the nerve fiber (**Figure 1C**).

**Figure 2.**
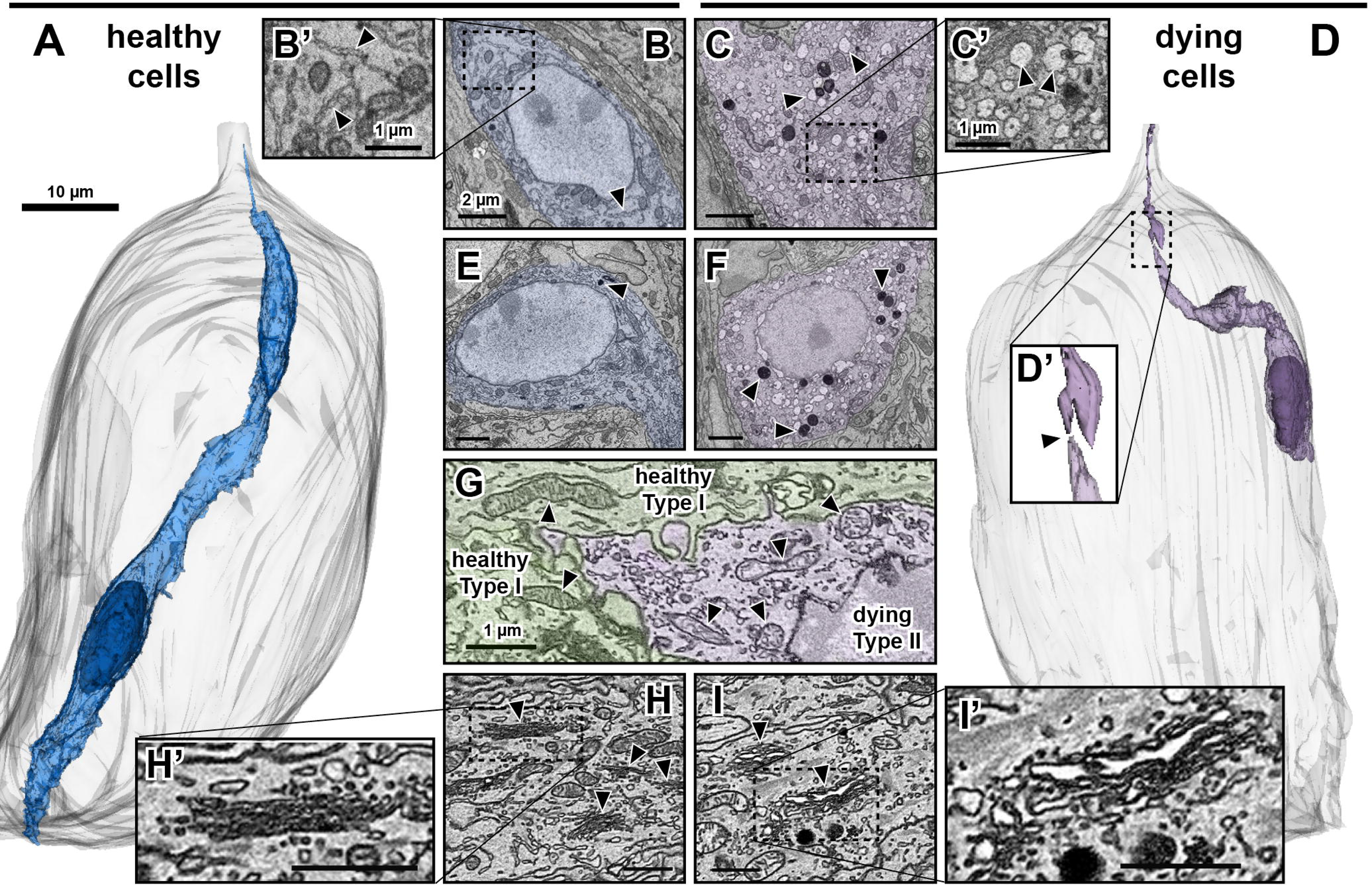
Morphological features of healthy and dying taste cells. **A.** Reconstruction of a healthy, mature Type II cell (blue) in the taste bud shell (gray). Scale bar is 10 µm. **B.** Micrograph of a healthy Type II cell, with arrows indicating endoplasmic reticulum (ER). Scale bar is 2 µm and applies to **B, C, E,** and **F**. **B’**. Enlarged region of healthy Type II cytosol showing normal ER (arrows). Scale bar is 1 µm. **C.** Micrograph of a dying Type II cell, with arrows indicating examples of swollen endoplasmic reticulum. **C’.** Enlarged region of dying Type II cell cytosol, featuring swollen endoplasmic reticulum. Scale bar is 1 µm. **D.** Reconstruction of a dying cell of unknown type (purple) in the taste bud shell (gray). Scale is the same as in **A**. **D’.** Enlarged region of dying taste cell reconstruction in **D**. Arrow indicates the point of separation between the main dying cell and its apical fragment. **E.** Micrograph of healthy Type II cell, with an arrow indicating the only visible lysosome in this cell profile. **F.** Micrograph of a dying Type II cell, with arrows indicating some of the numerous lysosomes apparent in the cell cytosol. **G.** Micrograph depicting two healthy Type I cells (green, left) and a dying Type II cell (purple, right). Arrows indicate mitochondria in healthy cells (left 2) and those of the dying cell (right 4). Scale bar is 1 µm and applies to **G, H, H’, I,** and **I’**. **H.** Micrograph of healthy cell cytosol with arrows indicating Golgi bodies. **H’.** Enlarged region of cytosol including a healthy Golgi apparatus. **I.** Micrograph of a dying Type II cell with arrows indicating swollen Golgi bodies. **I’.** Enlarged region of the swollen Golgi apparatus of a dying cell.

### Identifying lysosomes

The morphology of lysosomes can vary greatly over time and location in a cell (e.g., Dr. Jastrow’s electron microscopic atlas, http://www.drjastrow.de/WAI/EM/EMAtlas.html.) Taste cell lysosomes appear to fall into two categories. The first category contains roughly spherical membrane-bound vesicles of about 150-600 nm in diameter containing packed, electron dense granules. The second category contains larger, irregularly shaped structures up to 1-2 µm in the longest dimension. These larger lysosomes have inclusions of electron dense material embedded in a matrix comprising a combination of somewhat electron dense material and electron lucent material. We segmented profiles of lysosomes in every section of selected cells, being careful not to include any tangential or cross-sections of mitochondria which may have a similar appearance in single sections. Counts of lysosomes were performed using images taken from 3D reconstructions of cells primarily from dataset TF21.

### Estimating ER distention

To estimate the size of dying cell endoplasmic reticulum regions as compared to those of healthy cells, we chose three healthy and three dying cells at random. In these cells, we randomly chose among sections containing the cell nucleus, using an online random number generator to select the section number, and used Photoshop (Adobe) to measure 5 separate sections of ER at their widest points from each cell. These values were then averaged for our estimates.

### Nuclear Reconstructions and Display

Nuclear and taste bud perimeter traces were exported from the Reconstruct series files into MATLAB (Mathworks, Natick, MA) using custom MATLAB scripts (https://github.com/salcedoe/Dying_Taste_Cell_analysis). These traces were exported as a series of X, Y, and Z geometric vertices, which were organized in a table and sorted by cell identity and cell type. Vertices from individual nuclei were bound into 3D polyhedral meshes using the alphaShape function and visualized by plotting as 3D surface polygons.

### Lysosome and Cell Volume Analysis

Cell and lysosomal traces were imported as geometric vertices into MATLAB as described for the nuclear reconstructions. The vertices were then grouped by cell identity and cell type. To determine lysosome size, vertices from lysosome traces were converted into Point Cloud objects using the pointCloud tool from the MATLAB Computer Vision Toolbox. These point clouds were then segmented into distinct lysosome clusters based on a set Euclidean distance using the MATLAB pcsegdist function. Individual lysosome volumes were calculated by converting the lysosome point cloud clusters into 3D polyhedral meshes using the MATLAB alphaShape function, which then calculated the encased volume of each mesh. All generated 3D polyhedral surface meshes were visually inspected for morphological accuracy. Surface defects (e.g. large holes in the surface) were repaired using Manifold plus (https://github.com/hjwdzh/ManifoldPlus) and Meshfix 2.1 (https://github.com/MarcoAttene/MeshFix-V2.1). Taste cell volumes were then calculated from these inspected surface meshes. All cell meshes can be found on the github repository in the Lysosome Analysis/cellMeshes folder.

### Statistical Analysis

MATLAB was used to calculate the statistics for the lysosome sizes. Comparisons between separate cell categories (i.e. healthy Type II cells, early stage dying Type II cells, healthy Type III cells, late stage dying cells, etc.) with regards to lysosome volumes were performed with Kruskal-Wallis and ANOVA tests, depending on whether datasets qualified as skewed by a Kolmogorov-Smirnov test. Comparisons between separate cell categories with regards to cell volumes were performed using estimation statistics on the median difference. (estimationstats.com; Ho et al., 2019). The results of these cell volume estimation statistics are presented in **Supplemental Figure 1.**

### Code/Software Accessibility

Code was generated for lysosome volumetric analysis and is readily available on GitHub (https://github.com/salcedoe/Dying_Taste_Cell_analysis).

## Results

### General features of dying taste cells and their nuclei

The taste cells that we identify as dying share several distinct morphological features that distinguish them from mature taste cells (**Figure 2**). In the cytoplasm of dying cells, abundant, swollen endoplasmic reticulum and large lysosomes are among the most readily apparent and common of these features. The endoplasmic reticulum of dying cells appears swollen in comparison to that of mature taste cells; the width (roughly perpendicular to the longitudinal axis) of ER segments in dying cells is ∼250nm, while in healthy cells, the ER width is ∼90nm (**Figure 2B, B’, C, C’**). Healthy cells contain relatively small lysosomes, with median individual lysosome volumes ranging from 0.005-0.06 µm^3^ per cell. In contrast, lysosomes in dying cells tend to be larger, with median lysosome volumes ranging from 0.004-0.12 µm per cell (**Figure 2E, F**, **Figure 4**). Lysosome volumes differed significantly between the six different groups—I, II, III, IV, early dying, and late dying [Kruskal-Wallis test, H(5, n=3904) = 328.2, p<0.0001]. When we directly compared the lysosomes from healthy and early dying Type II cells in a post-hoc, multiple comparison analysis, we found the lysosomes in the dying cells to be significantly larger than the lysosomes in the healthy cells (p<0.0001). Mitochondria in dying cells often display signs of degradation—instead of the stacked cristae of healthy mitochondria or the tubular cristae of atypical mitochondria, mitochondria in dying cells often contain irregular, sparse cristae (**Figure 2G**). Golgi bodies in dying cells, like the endoplasmic reticulum, appear swollen (**Figure 2I, I’**) when compared to those of healthy, mature taste cells (**Figure 2H, H’**). A subset of dying taste cells feature more obvious signs of apoptosis: cell shrinkage and even cell fragmentation into apoptotic bodies (**Figure 2A, D**). We deem these cells “late stage” dying cells, as opposed to the “early stage” dying cells, which are not fragmented and are still identifiable by cell type. Late stage dying cells are smaller than healthy cells, ranging in volume from 403-457 µm , while healthy Type II and III cells for which volume was measured range from 524-1172 µm . Interestingly, early stage dying Type II and III cells tend to be slightly larger than their healthy counterparts as well as late stage dying cells, ranging from 319-1984 µm^3^ (**Figure 3, Supplementary Figure 1**). Late stage dying cells are fragmented into multiple separate objects. We presume these objects to be fragmented apoptotic bodies when they are in proximity to the main cell body and mirror its cytosolic qualities (**Figure 2D**).

**Figure 3.**
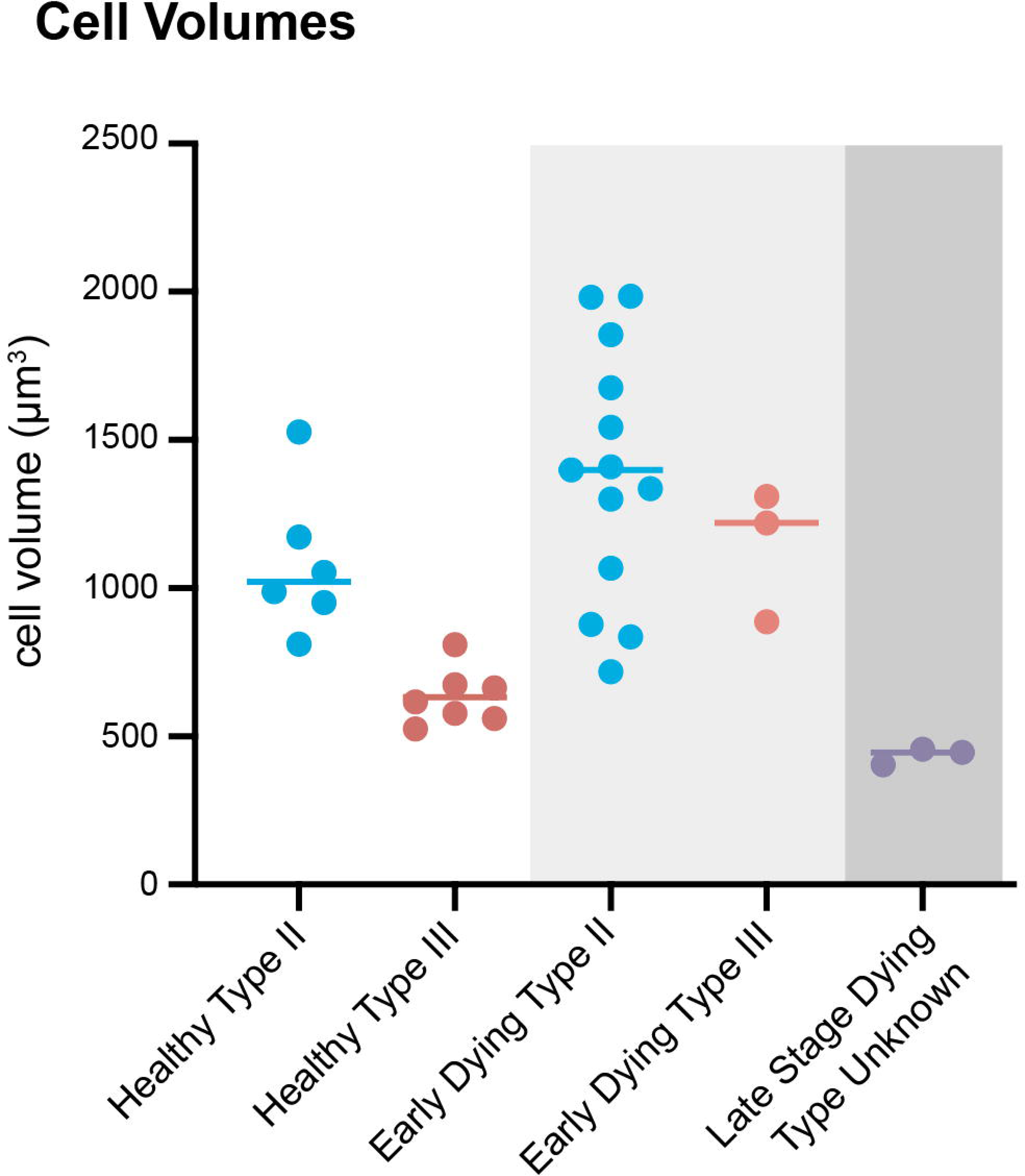
Cell volumes of taste cells. Graph depicts total cell volumes of healthy Type II and III cells (left), early dying Type II and III cells (middle, light gray), and late stage dying cells (right, dark gray).

**Figure 4.**
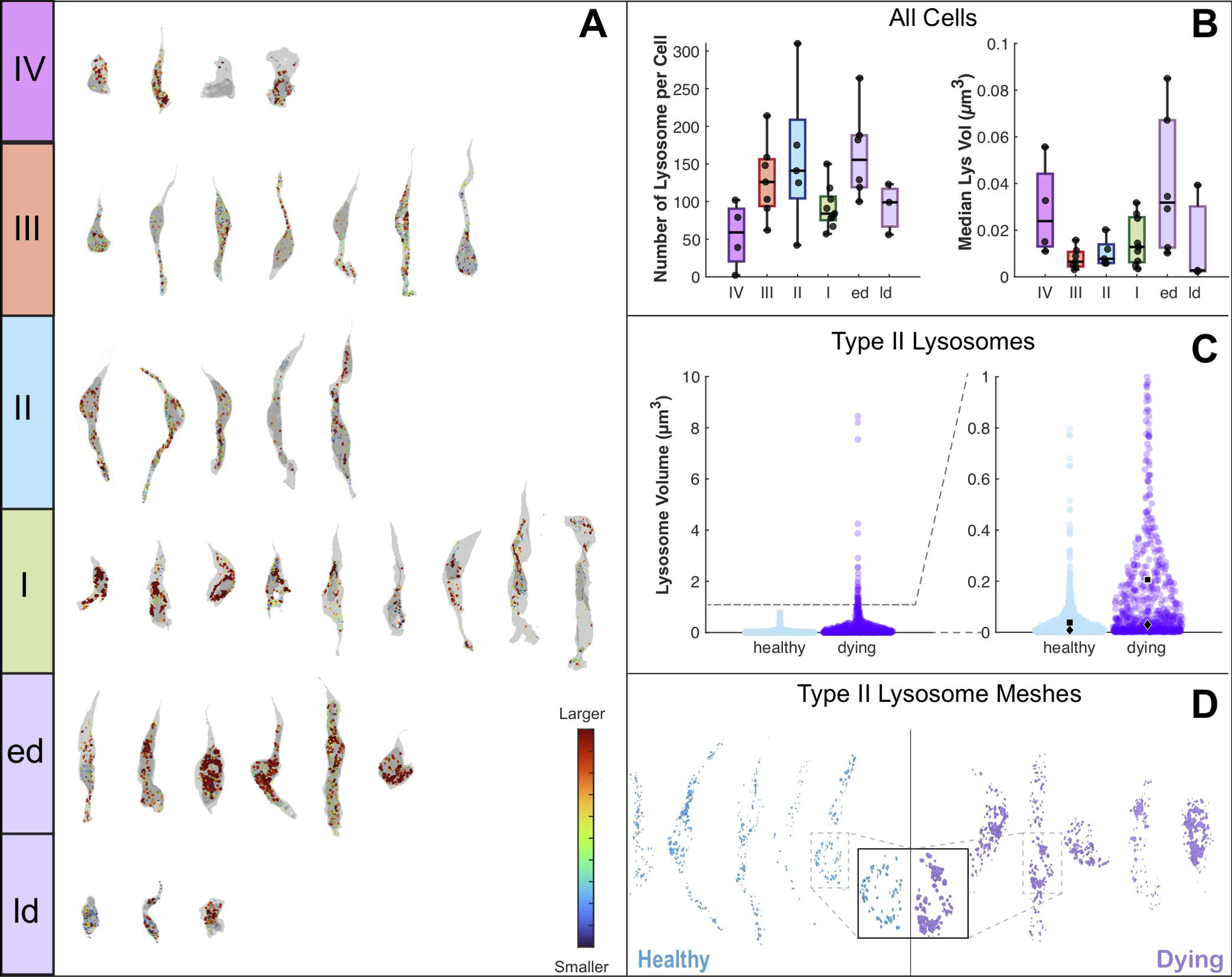
Lysosomes in taste cells. **A.** Reconstructions of selected taste cells (grey) and their lysosomes (color coded by size from small (blue) to large (red)). Reconstructions are divided into cell types: IV, III, II, I, early dying (**ed**), and late dying (**ld**). **B.** Lysosome count per cell (left) and lysosome median volume by cell (right). Each data point represents a single cell. Data is binned by cell type, as in **A. C.** Swarm charts of lysosome volumes of healthy Type II cells and early stage dying Type II cells showing the entire range of volumes (left) and a cropped view (right) of the volumes below 1µm . Black squares indicate mean volume, while diamonds indicate median volume. **D.** Reconstructions of the lysosome meshes from healthy Type II cells (blue) and an early stage dying Type II cells (purple). Inset shows an enlarged view of the perinuclear area from a healthy and dying cell.

Nuclei in dying cells likewise differ from those of mature taste cells (**Figure 5**). In dying cells that feature swollen endoplasmic reticulum and large lysosomes, the nuclear membranes exhibit clear, distended regions of separation between the inner and outer nuclear lamellae (**Figure 5B, C**). The degree of nuclear membrane separation varies among dying cells, with individual distended regions ranging from ∼160 to 500 nm between the inner and outer leaflets (**Figure 5B, C, E’, G’**). Since this intra-membrane region is contiguous with the inner endoplasmic reticular space (Lindenboim et al., 2020), the distended nuclear membrane of dying cells may be an extension of the swollen endoplasmic reticulum, or vice versa. In a small subset of dying cells, nuclei feature accumulations of dense heterochromatin, a common characteristic of apoptotic cells (D’Arcy, 2019; Snigirevskaya and Komissarchik, 2019) (**Figure 5E’, G’**). In one cell that appears to have fragmented into multiple apoptotic bodies, the nucleus is likewise fragmented into multiple bodies, which is consistent with caspase-induced breakdown of nuclear lamins (for review, Fink and Cookson, 2005) (**Figure 5E, E’**). Other nuclei in late stage dying cells possess large invaginations, which are perhaps harbingers to nuclear fragmentation (**Figure 5G, G’**).

**Figure 5.**
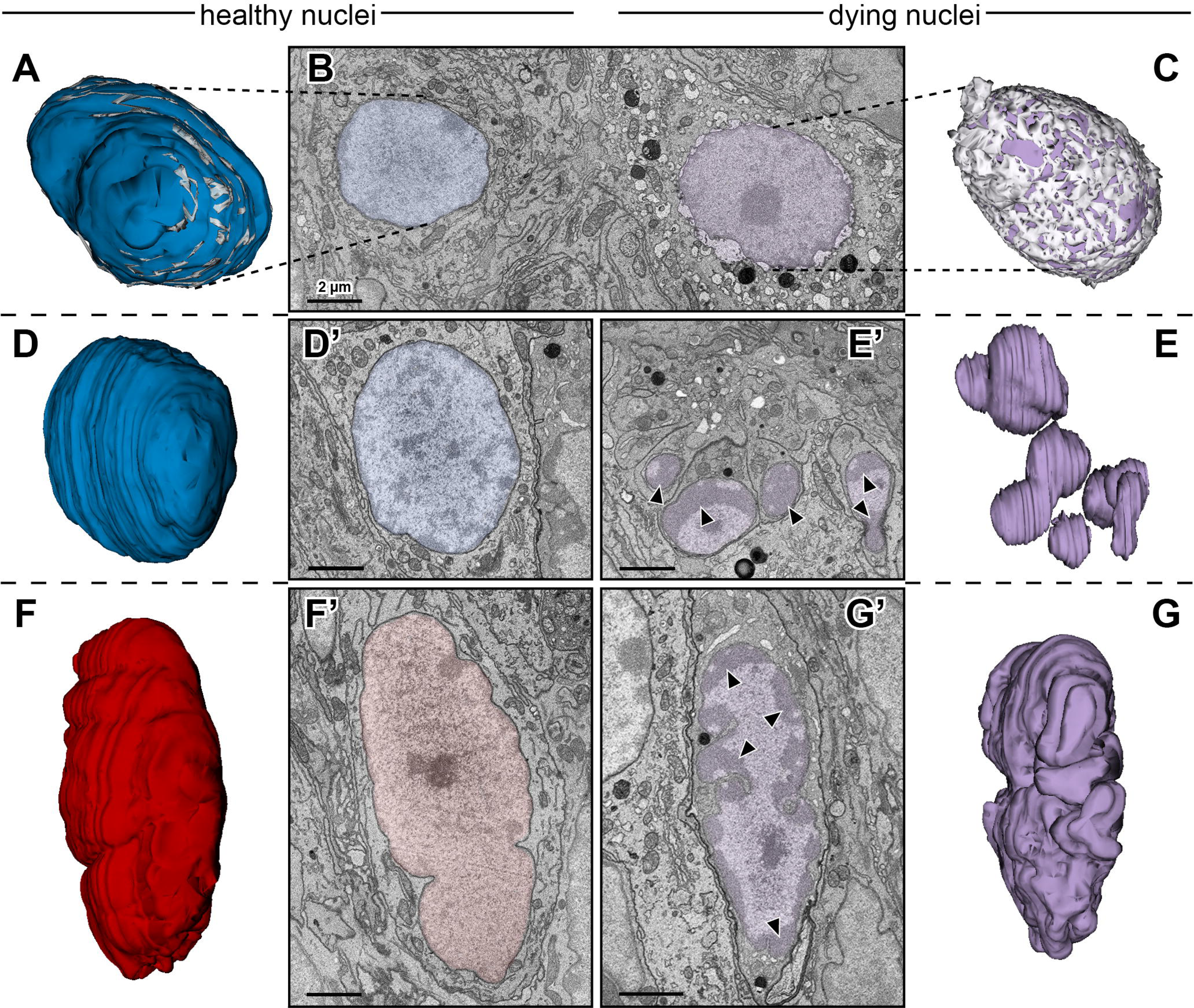
Morphological features of healthy and dying taste cell nuclei. **A.** Reconstruction of healthy Type II cell nucleus (blue) with distended regions between inner and outer nuclear membranes reconstructed in gray. **B.** Micrograph of a healthy Type II cell nucleus (that reconstructed in **A**) (blue, left) and a nucleus of a dying Type II cell on the right (purple) showing distended regions between the inner and outer nuclear membrane. Scale bar is 2 µm, and applies to all micrographs in this figure. **C.** Reconstruction of the nucleus shown in **B**, showing in the inner nuclear region in purple, and the distended regions between the inner nuclear membrane in lighter purple. Of the dying cell nuclei, this is the only nucleus for which these distended regions were segmented and reconstructed. Thus, the reconstructions in **E** and **G** do not feature equivalent reconstructions of the distended nuclear membrane regions**. D, D’.** Reconstruction (**D**) and micrograph (**D’**) of healthy Type II cell nucleus. **E, E’.** Reconstruction (**E**) and micrograph (**E’**) of a fragmented nucleus of a dying cell of unknown type. Arrows indicate regions of heterochromatin expansion. **F, F’.** Reconstruction (**F**) and micrograph (**F’**) of a healthy Type III cell nucleus. **G, G’.** Reconstruction (**G**) and micrograph (**G’**) of the shrunken nucleus of a dying cell of unknown type. Arrows indicate regions of heterochromatin expansion.

### Synapses in dying cells

The genesis and degeneration of taste cells necessitates remodeling of synaptic contacts and nerve fibers. To ascertain whether dying cells might still be communicating to taste nerves, we investigated possible synapses between dying cells and afferent nerve fibers. Late stage dying cells did not exhibit structures consistent with synapses onto afferent nerve fibers. At points of contact with nerve fibers, late stage dying cells lacked both the atypical mitochondria that characterize Type II cell synapses onto nerve fibers as well as the pre-synaptic clusters of vesicles characteristic of Type III cell synapses (**Figure 6A**).

**Figure 6.**
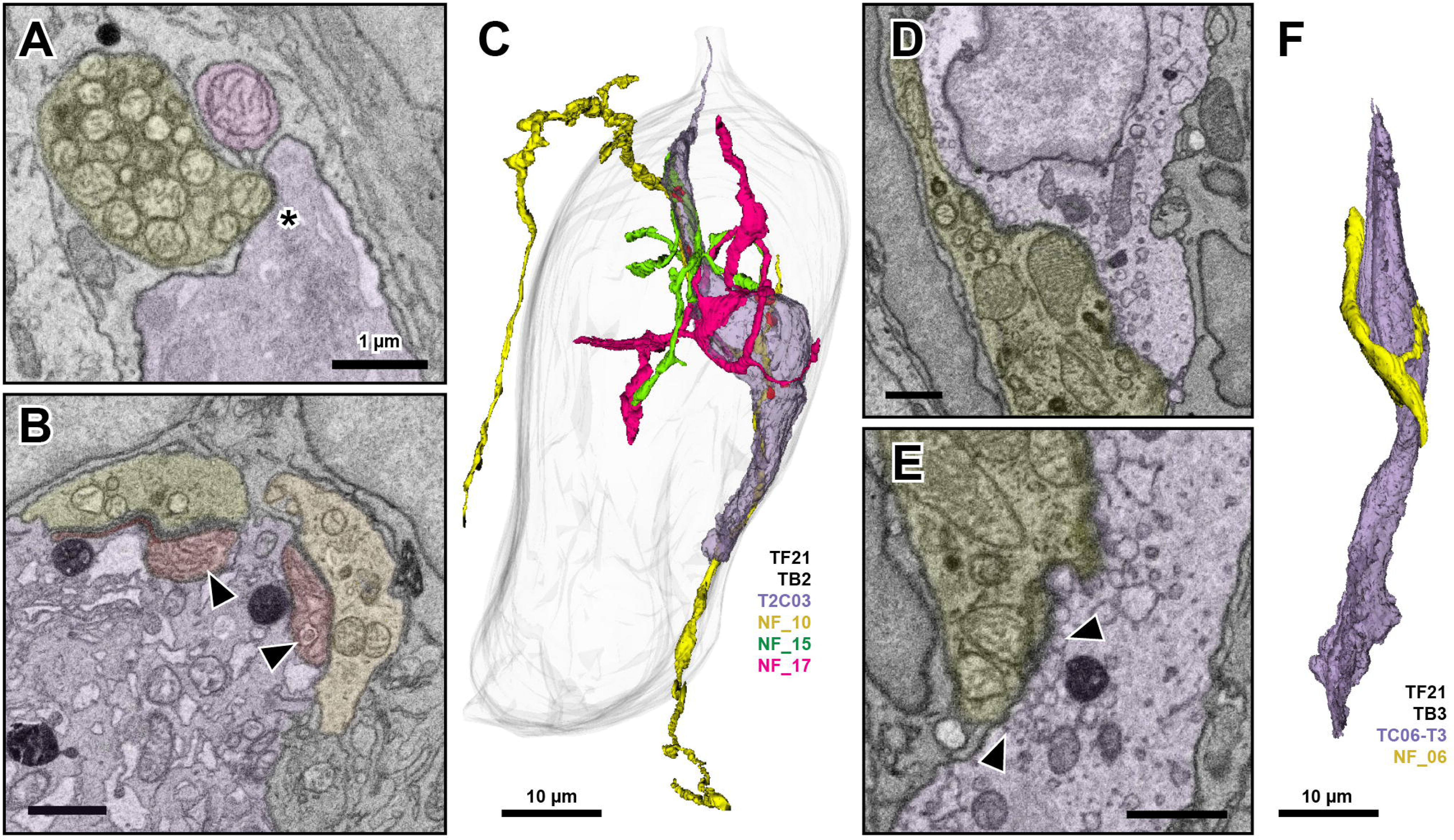
Synapses in dying cells. **A.** Micrograph of the point of contact between a late stage dying cell of unknown type (purple) and an adjacent nerve fiber (yellow). Asterisk indicates region of contact. Scale bar is 1 µm and applies to all micrographs in this figure. **B.** Synaptic sites between two nerve fibers (yellow-green and yellow) and a dying Type II cell (purple). Arrows indicate atypical mitochondria (red), which feature abnormal cristae patterns. **C.** Reconstruction of a dying Type II cell (purple) and the three nerve fibers that receive synapses from it in the taste bud shell (gray). Atypical mitochondria are red. Only the orange fiber exits the base of the taste bud. The green and pink fibers appear to be nerve fiber fragments. Scale bar 10 µm. **D.** Micrograph showing a region of contact between a dying Type III cell (purple) and the adjacent nerve fiber (yellow). **E.** Enlarged micrograph showing a different point of contact between the dying Type III cell (purple) and the nerve fiber (yellow) in **D**. Arrows indicate large, swollen structures (upper arrow) and typical synaptic vesicle structures (lower arrow) at the point of contact with the nerve fiber. **F.** Reconstruction of a dying Type III cell (purple) and the nerve fiber it borders. This nerve fiber is a fragment and does not exit the base of the bud.

In contrast, most early stage dying cells were Type II cells still showing the presence of atypical mitochondria characteristic of synaptic contacts from this cell type. These atypical mitochondria appeared at sites of contact with nerve fibers, but these nerve fibers did not always exit the taste bud, suggesting nerve fiber fragmentation. This observation suggests that early stage dying Type II cells retain gross synaptic structures, but may not always be capable of signaling information to the CNS since the nerve fiber has fragmented (**Figure 6B, C**) and is no longer connected to the CNS. The atypical mitochondria in early stage dying Type II cells feature irregular, “loose” cristae (**Figure 6B**) when compared to the tubular cristae of atypical mitochondria in mature, healthy Type II cells (**Figure 1B**).

A minority of the early stage dying cells are Type III cells. These cells share similar cytoplasmic quality with early stage dying Type II cells, but lack atypical mitochondria. Instead, at points of contact with nerve fibers, we observe clear, membrane enclosed profiles that are larger than typical synaptic vesicles. Synaptic vesicles in healthy Type III cells range from 40-60 nm (Yang et al., 2020); membrane-bound objects at the site of contact between a dying Type III cell and a bordering nerve fiber range from ∼30-300 nm. We tentatively mark these structures as degrading synaptic contacts, although the occurrence of signal transmission at these sites is unknowable with our current methods (**Figure 6D, E**). As with some fibers innervating early stage dying Type II cells, some nerve fibers contacting the putative early stage dying Type III cells do not exit the bud, suggesting nerve terminal fragmentation (**Figure 6F**). In TF21_ TB2, from which we have a complete connectome (Wilson et al., 2022), 4 of 7 (57%) nerve fibers innervating dying cells are fragments that do not appear to exit the bud, while just 8 of 25 (32%) of nerve fibers innervating healthy cells appear to be fragments.

### Type I cells engulf dying cells

Apoptotic cells elsewhere in the body are generally phagocytosed by elements of the immune system, often prior to the development of apoptotic bodies (D’Arcy, 2019). In the nasal epithelium, which is largely devoid of immune cells, sustentacular cells phagocytose neighboring epithelial cells (Suzuki et al., 1996B). Type I cells are considered the glial-like cells of the taste bud (for review, Chaudhari and Roper 2010). Type I cells possess diaphanous processes that extend around and between neighboring taste cells and are well positioned to engulf dying cells and any apoptotic bodies they might generate. Indeed, we observe signs of Type I cells engulfing dying cells and their fragmented apoptotic bodies (**Figure 7A-C**). In addition, membrane bound objects matching the cytosolic presentation of a dying cell are often seen in the cytoplasm of immediately adjacent Type I cells (**Figure 7C, E**). In regions of Type I cells that border dying cells, lysosomes tend to be larger, much like the lysosomes in the dying cells themselves (**Figure 4**, **Figure 7D, D’’**). The 3 largest individual lysosomes in Type I cells range from 7.5-9.1 µm , which is several times the volume of the median volume lysosomes in either healthy or dying cells. In one case, a Type I cell that neighbors an early stage dying cell appears to have engulfed large, vacuous, membrane enclosed bodies. These bodies closely resemble the cytosolic appearance of the dying cell rather than the Type I cell itself. Thus, we presume this material originated in the dying cell (**Figure 7E**). We have never observed either immune cells or other non-taste cells within the confines of the taste bud that could be involved in the elimination of dying cells.

**Figure 7.**
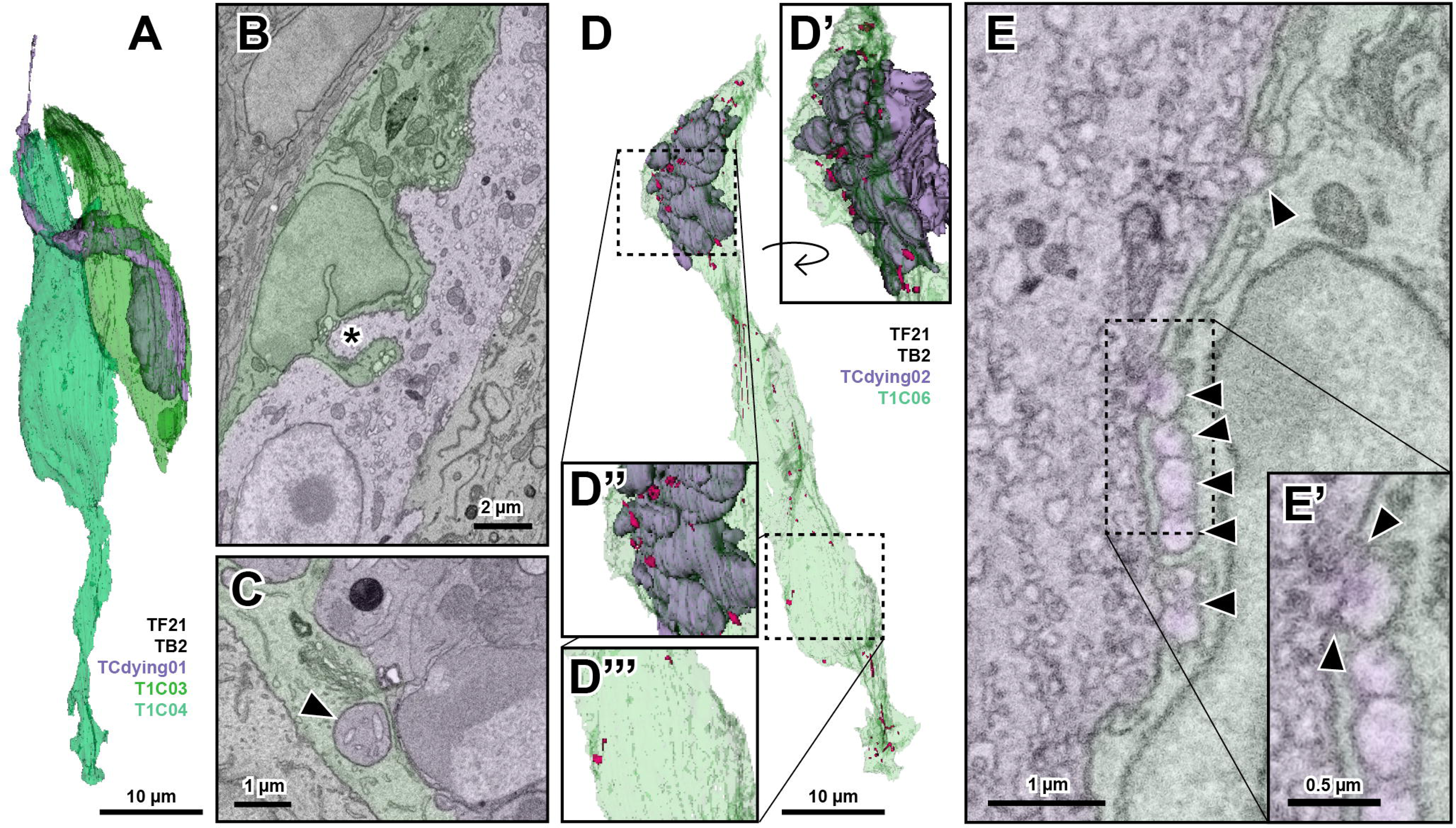
Type I cells as they relate to dying cells. **A.** Reconstruction of a late stage dying cell (purple) and the two Type I cells (green) that surround it. **B.** Micrograph of an early stage dying Type II cell (purple) and a neighboring Type I cell (green). Asterisk indicates a region of the dying cell nearly surrounded by the neighboring Type I cell, likely an area of phagocytosis. **C.** Micrograph of a late stage dying cell (purple) and the bordering Type I cell (green). Arrow indicates a presumed apoptotic body, which is completely surrounded by the Type I cell. **D.** Reconstruction of a late stage dying cell (purple) and the neighboring Type I cell (green). Lysosomes in the Type I cell are depicted in pink. **D’.** Rotated, enlarged view of the Type I cell partially surrounding the dying cell. **D’’.** Enlarged region of D which highlights the lysosomes (pink) in the region of the Type I cell that borders the dying cell. **D’’’.** Enlarged region of D which highlights the lysosomes (pink) in the region of the Type I cell that does not border the dying cell. **E.** Micrograph of a Type I cell (green) that borders an early dying cell (purple). Arrows indicate large, membrane enclosed profiles that the Type I cell may have engulfed from the dying cell. **E’.** Enlarged region of E which highlights the membrane continuity between the dying cell and one such profile that is perhaps in the process of being engulfed by the neighboring Type I cell.

### Dying taste cells in the context of the bud

In total, we identify 21 dying cells in 5 different taste buds over two separate datasets. In the two taste buds wholly contained within the segmented block of tissue, 4 out of 84 and 5 out of 86 total taste cells appear to be dying according to our previously discussed morphological criteria. Of these dying cells, early stage dying cells are still identifiable as belonging to a mature taste cell type. These cells feature some, but not all, characteristics of dying cells. They tend to display swollen endoplasmic reticulum, degrading mitochondria, blebby nuclear membranes, swollen Golgi bodies, and large lysosomes. Of these early stage cells, most are Type II cells that maintain aspects of Type II cell morphology and atypical mitochondria. The remaining early stage dying cells share qualities with mature Type III cells. No early stage dying cells exhibit Type I cell morphology although Type I cells are the majority of the cells in a taste bud. Late stage dying cells could not be identified as to taste cell subtype, because of the more advanced signs of cell degradation: cell fragmentation, nuclear fragmentation, and notable reorganization of dense regions of heterochromatin in the nucleus. We thus label these cells as “unknown type” (**Figure 8A**, **Figure 9**).

**Figure 8.**
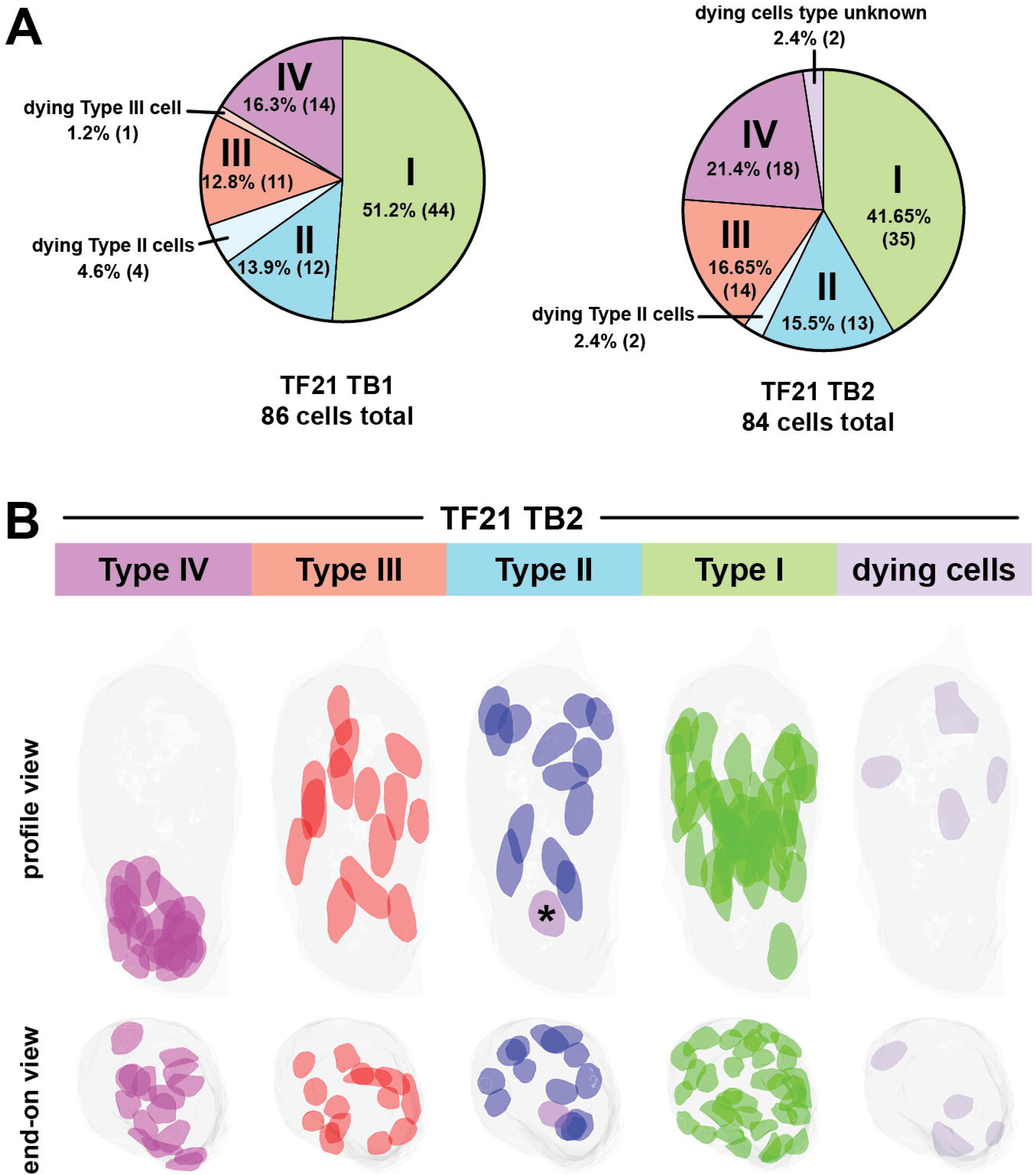
Dying cells in the context of the bud. **A.** Pie charts depicting the cell type composition of two taste buds: TF21 TB1 and TF21 TB2. These two taste buds are the only taste buds that are fully contained in the boundaries of the tissue block. **B.** Diagram depicting the locations of cell nuclei within the taste bud TF21 TB2 according to cell type. Immature Type IV cell nuclei (pink-purple) are on the left, followed by Type III cell nuclei (red), Type II cell nuclei (blue), Type I cell nuclei (green) and dying cell nuclei (dusty purple). One nucleus in the Type II cell panel is labeled as purple (asterisk), because the cell it inhabits is immature and we cannot discern whether it would have developed into a Type II or Type III cell. The taste bud and nuclei are shown in profile (top row) and from a top down view of the taste bud (bottom).

**Figure 9.**
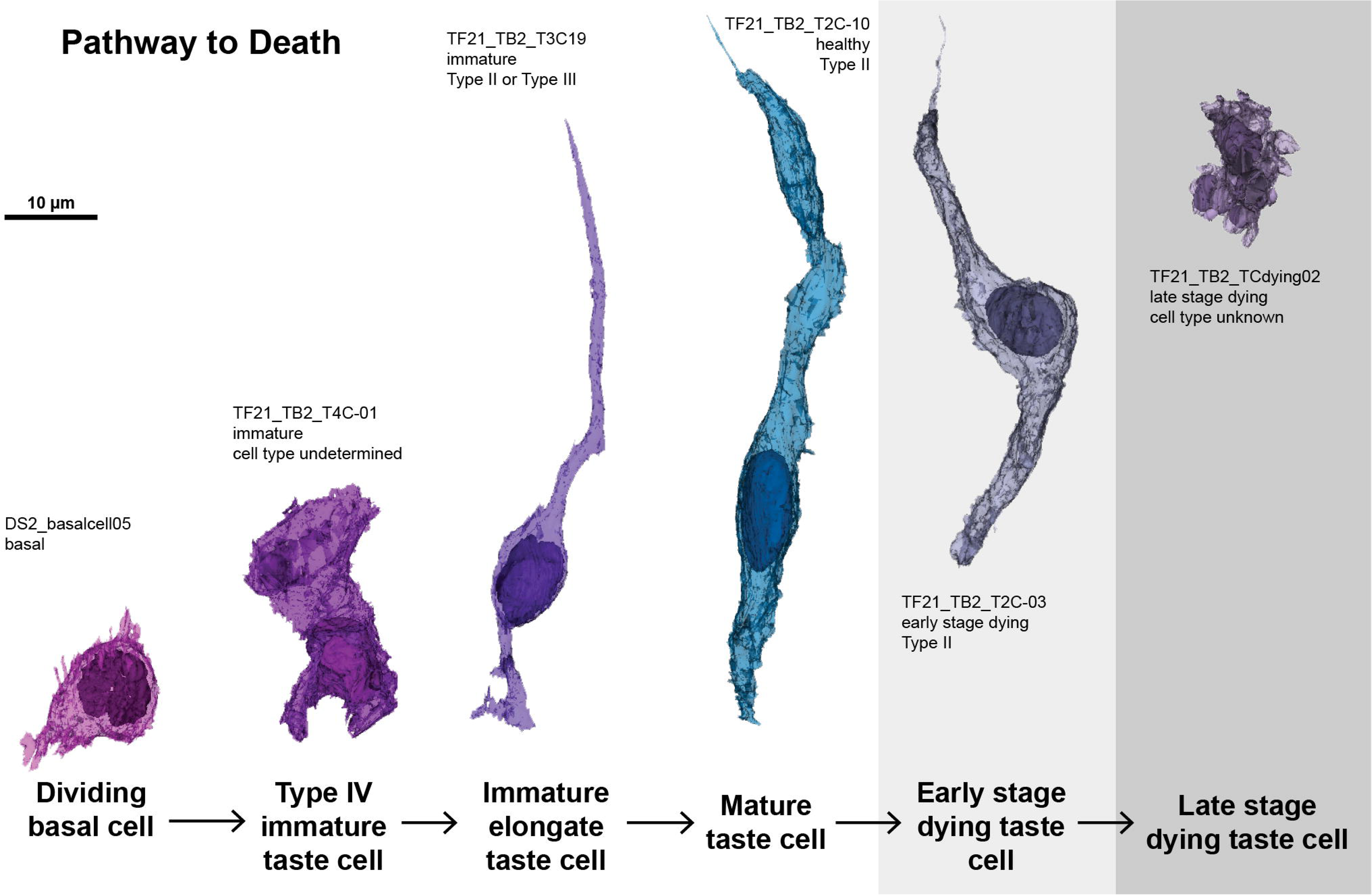
Pathway to death. Reconstructions of cells summarizing the pathway from birth to death of a taste cell. From left to right: dividing basal cell, Type IV immature taste cell, Immature Type II or III taste cell, Type II mature taste cell, Type II early stage dying cell, Late stage dying cell of unknown type.

### Location of dying cells within the taste bud

Post-mitotic, immature Type IV taste cells are known to enter the taste bud near the basilar membrane and inhabit the bottom 1/3 of the bud (Barlow, 2015; Yang et al., 2020). As they mature, taste cells extend into the upper portions of the bud, eventually reaching the taste pore. We hypothesized that dying cells would be restricted to the upper regions of the taste bud, farther away from their origins in the basal regions of the bud consistent with a general migration from basal to apical within the epithelium. Indeed, the nuclei of dying cells tend to be in the top 1/3 of the taste bud, although taste cells at the upper regions of the bud do not all display hallmarks of degeneration (**Figure 8B**).

## Discussion

The data we present suggest an apoptotic pathway for the death of Type II and III taste receptor cells (**Figure 9**), but not for glial-like Type I cells. As the receptor cells progress towards death, the ER and Golgi swell, lysosomes enlarge, mitochondrial cristae become disorganized, and the inner and outer leaflets of the nuclear membrane separate (**Figures 2, 4, and 5**). These features are consistent with apoptosis (D’Arcy, 2019; Snigirevskaya and Komissarchik, 2019), and agree with reports of apoptotic markers within taste buds (Zeng and Oakley, 1999; Zeng et al., 2000; Huang and Lu, 2001; Takeda et al., 1996). Early stage dying cells tend to be larger than healthy or late stage cells, indicating that cells swell slightly before late stage cell death (**Figure 3**). As cells progress to the late dying stage, heterochromatin reorganization becomes more pronounced, cell volume reduces, and apoptotic bodies separate from the main cell body (**Figures 2, 3, and 5**).

We conclude that the dying cells we describe are undergoing apoptosis. If taste cells were dying by non-apoptotic methods, we would expect different morphologies. Cells undergoing necrosis would manifest substantial cell swelling, extracellular spillage of cell contents, and infiltration of immune cells (D’Arcy, 2019; Lakshmanan et al., 2022). Instead, we observe cell shrinkage in late stage dying cells, formation of putative apoptotic bodies, and no evidence of immune cell infiltration. Interestingly, early stage dying cells are slightly larger than healthy cells (**Figure 3, Supplementary Figure 1**). That cells swell before shrinking is not necessarily inconsistent with apoptosis. Reduced ATP production on account of degrading mitochondria might disrupt the Na /K -ATPase, which can result in swelling (Chen et al., 2014). In autophagy, dying cells would form autophagosomes, which sequester and degrade macroproteins and organelles (Eskelinen et al., 2011). We do not observe such structures in dying Type II or III cells, although autophagosome-like structures occasionally appear in Type I cells. However, we cannot determine if these structures indicate cell death or repair.

In the two complete buds contained within the samples, few cells per bud appear to be undergoing apoptosis: 4 of 84 (4.7%) and 5 of 86 (5.8%). Some previous studies estimate the dying cell population at 1-3 dying cells per bud (Ueda et al., 2008; Huang and Lu, 2001; Takeda et al., 1996), while others estimate it at ∼9 cells per bud (Zeng and Oakley, 1999; Zeng et al., 2000) (**Table 1**). Our observed prevalence of apoptotic cells may be higher than the lower estimates due to the relatively short period (1-3 hrs) during which an apoptotic cell is positive for classical apoptotic markers, i.e. TUNEL and ssDNA (Gavrieli et al., 1992). The studies that estimated a larger dying cohort used p53 and Caspase-2 as markers, both of which are initiating apoptotic factors whose presence likely precedes the morphological changes we report (Zeng and Oakley 1999; Zeng et al., 2000, Bouchier-Hayes and Green, 2012). Beidler and Smallman (1965) reported that rat taste buds lose approximately half their cells every 10 days. If mouse taste buds are similar, we expect 4-5 dying cells per day in a bud of 80-100 cells. Our results are consistent with this estimate.

**Table 1.** Reported numbers of dying cells in taste buds.

| Citation | Animal model | Death identification method | Observed dying cells per section | If bud = 40 μm across, dying cells/bud |
| --- | --- | --- | --- | --- |
| Ueda et al 2008 | Rat | ssDNA stain | 1.2 cells / 16 μm section | ~3 |
| Zeng & Oakley 1999 | Mouse | p53 (early death signal) | 11.4% | 11.4% (~9 cells) |
| Zeng et al 2000 | Mouse | Caspase-2 (protease) | 11% +/- 0.8% | 11% (~9 cells) |
| Huang & Lu 2001 | Guinea pig | Tunel stain | ~1-2 tunel+ cells / 20 μm section | 2-4 |
| Takeda et al 1996 | Mouse | Tunel stain | ~0.14 tunel+ cells / 6 μm section | 0.93 |

Of early stage dying taste cells, the majority are Type II cells and a few are Type III cells—we observe no dying Type I cells. Buds contain fewer Type III than Type II cells, and Type III cells are the longest-lived, making it less likely that we would observe them dying in a single snapshot of a dynamic organ. Surprisingly, we do not observe apoptotic Type I cells. Type I cells are the most common cell type (∼50% of all cells; Yang et al., 2020), and are described as the shortest-lived (Farbman, 1980; Perea-Martinez et al., 2013; Gross et al., 2017; Yang et al., 2020). If a bud contains ∼100 cells, we expect ∼50 to be Type I cells. If, as according to the estimates of Beidler and Smallman (1965) and Yang et al (2020), half of this population is renewed every 10 days, then we would expect ∼2.5 Type I cells dying per day in each taste bud. Since this estimate is based on the entire taste cell population, and Type I cells reportedly have the shortest lifespan (Farbman, 1980; Hamamichi et al., 2006; Perea-Martinez et al., 2013; Gross et al., 2017), the estimate is likely conservative. However, we do not observe apoptotic Type I cells.

What then, is their ultimate fate? Ueda and colleagues reported a population of apoptotic cells interpreted as Type I cells in rats (Ueda et al., 2008). The marker used to identify these cells (human blood antigen H) is, however, not specific to Type I cells (Ueda et al., 2003). Other estimates of Type I cell lifespans are based on perdurance of nucleotide labeling during progenitor cell division; Farbman (1980) estimated “dark” cells (an earlier term apparently equivalent to Type I cells) to have a lifespan of ∼7 days, and Perea-Martinez (2013) and colleagues found cells lacking Type II or III markers (presumed Type I cells) to have a ∼16 day lifespan. If Type I cells are exiting the bud rather than dying, previous studies would not necessarily capture that process.

Conceivably, Type I cells may exit the taste bud apically or laterally, to be eventually sloughed off the lingual epithelium with non-taste epithelial cells. Alternatively, Type I cells may be dying too quickly to capture in our sampling paradigm, or may die via an alternative cell death process which shows no obvious morphological features. We do occasionally observe autophagosome-like profiles in Type I cells, but cannot determine whether these structures indicate cell death, cell repair, or phagocytosis of neighboring dying cells. Possibly, the Type I cells undergo autophagy following phagocytosis of other dying cells. The cells we describe as late stage dying cells could include Type I cells, but are more rare than expected. A final alternative is that Type I cells are more long-lived than previously estimated, and that Type II and III cells turn over while the Type I cell population remains relatively constant. Our data do not resolve how and whether Type I cells die or otherwise exit the taste bud.

Early stage dying Type II and III cells feature synaptic structures in various stages of degradation, suggesting impaired signaling to nerves. Moreover, these dying cells often contact postsynaptic nerve fibers that, seemingly, do not exit the bud and thus cannot signal to the CNS. In the face of wholesale turnover of taste receptor cells, taste nerves are constantly remodeling (Whiddon et al., 2023). Large “end-bulbs” are associated with nerve branch retraction (Zaidi et al., 2016; Whiddon et al., 2023) and we occasionally observe enlarged nerve processes near late stage dying cells (**Figure 6A**). More often, synapses from dying cells contact nerve fiber fragments (**Figure 6B-F**). Considering these data, we hypothesize that nerve fiber fragments that receive synapses from early stage dying cells become separated from their trunk nerve fibers as a part of the remodeling process. Such nerve fragments are not reported by Whiddon et al (2023), who rely on sparse GFP label to follow individual fibers within a taste bud. A possible explanation for this discrepancy is that once a nerve fiber fragment detaches from the nerve trunk, it’s ability to maintain pH homeostasis degrades, which would cause cytoplasmic acidification and would likely quench the GFP fluorescence (Roberts et al., 2016). We cannot, however, rule out the possibility that these nerve fiber fragments are attached to fibers exiting the bud, but that our data do not allow us to accurately trace the full nerve profile. In our dataset, it can be challenging to follow processes with diameters less than 70 nm (the section thickness) as they wend their way between other cell and fiber processes.

Type I cells play a crucial physiological role in taste buds, analogous to the role of astrocytes in regulating neural transmission at tripartite synapses in the brain (Araque et al., 1999; Dando et al., 2012; Kataoka et al., 2012; Rodriguez et al., 2021). Our data present a novel role for glial-like Type I cells in the homeostasis of taste bud structure, i.e. Type I cells engulf and remove neighboring apoptotic cells. In many tissues, immune cells fill this phagocytic role (D’Arcy, 2019) but we find no evidence for immune cells within taste buds. Type I cells, considered the glial-like support cells of the taste bud, are well poised for a phagocytic role. Type I cells most directly resemble astrocytes of the CNS—they degrade excess ATP, a taste neurotransmitter, and appear to sense and facilitate signaling at taste synapses (Roper, 2013; Rodriguez et al., 2021). While not the primary phagocytic glia in the CNS, astrocytes phagocytose and degrade synapses, distal dendrites of injured neurons, and dying microglia (Morizawa et al., 2017; Kono et al., 2021; Zhou et al., 2022). In the olfactory system, glial-like Olfactory Ensheathing Cells regularly phagocytose and degrade dying olfactory neurons which, like taste buds, make up a sensory cell population that is renewed repeatedly (Nazareth et al., 2021). Precedence also exists for supporting cells in sensory epithelia engulfing dying cells, as in the olfactory epithelium (Suzuki et al., 1996B). Previous studies found Type I taste cells containing large, dense bodies within their cytosol (Takeda et al., 1996; Farbman, 1969; Fujimoto and Murray 1970; Olivieri-Sangiacomo, 1970; Farbman, 1985). Our data are consistent with these observations, leading us to the hypothesis that Type I cells phagocytose and degrade debris from apoptotic Type II and III cells.

Our data point to apoptosis as the major mechanism of death for murine taste cells. The dying cells we identify feature several hallmarks of apoptosis, and can be categorized into either early or late stages of death. Those that retain synaptic contacts with nerve fibers during early stages of cell death are likely impaired in their ability to communicate to the CNS, as many of their target nerve fibers are fragmented. Type I cells appear to engulf dying cells, identifying a novel, phagocytic role for Type I taste cells. We do not, however, identify morphologically recognizable Type I cells in the process of cell death. Whether, how, and when Type I cells die or exit the taste bud remains unclear.

**Supplementary Figure 1. Estimation statistics for the median difference of cell volumes between cell categories.** The median differences between healthy and late stage dying cells (**A**); early stage and late stage dying cells (**B**); healthy Type II and early stage dying Type II cells (**C**); and healthy Type III and early stage dying Type III cells (**D**) as shown in Gardner-Altman estimation plots generated using estimationstats.com (Ho et al., 2019). For each comparison, both groups are plotted on the left axes; the mean difference is plotted on a floating axes on the right as a bootstrap sampling distribution. The mean difference is depicted as a dot; the ends of the vertical error bar (black) indicate the 95% confidence interval. All calculated confidence intervals indicate that the compared groups are different.

